# Rhythmic clock gene expression in Atlantic salmon parr brain

**DOI:** 10.1101/2021.08.18.456820

**Authors:** Charlotte M Bolton, Michaël Bekaert, Mariann Eilertsen, Jon Vidar Helvik, Herve Migaud

## Abstract

To better understand the complexity of clock genes in salmonids, a taxon with an additional whole genome duplication, an analysis was performed to identify and classify gene family members (*clock, arntl, period, cryptochrome, nr1d, ror*, and *csnk1*). The majority of clock genes, in zebrafish and Northern pike, appeared to be duplicated. In comparison to the 29 clock genes described in zebrafish, 48 clock genes were discovered in salmonid species. There was also evidence of species-specific reciprocal gene losses conserved to the *Oncorhynchus* sister clade. From the six period genes identified three were highly significantly rhythmic, and circadian in their expression patterns (*per1a*.*1, per1a*.*2, per1b*) and two was significantly rhythmically expressed (*per2a, per2b*). The transcriptomic study of juvenile Atlantic salmon (parr) brain tissues confirmed gene identification and revealed that there were 2,864 rhythmically expressed genes (*p <* 0.001), including 1,215 genes with a circadian expression pattern, of which 11 were clock genes. The majority of circadian expressed genes peaked two hours before and after daylight. These findings provide a foundation for further research into the function of clock genes circadian rhythmicity and the role of an enriched number of clock genes relating to seasonal driven life history in salmonids.

## 1 INTRODUCTION

The importance of biological time keeping is apparent across all organisms, from bacteria to humans (Wulund and Reddy, 2015). These time-dependent adaptations can last seconds or minutes and recur throughout the day (ultradian) or endure days or months (infradian). They are an evolutionary trait which enables all living organisms to maximise fitness in anticipation of endogenous (molecular and cellular) and external stimuli or zeitgebers (Andreani et al., 2015; Sánchez-Vázquez et al., 2021). Circadian rhythms are endogenous oscillatory expression of genes with a periodicity of approximately 24-hours. These rhythms are the expression of endogenous clocks which synchronises biochemical, physiological, and behavioural responses enabling organisms to respond to diel environmental changes (Li et al., 2015). The transcription of 43% of all protein-coding genes in mice displayed circadian rhythms across numerous organs (Zhang et al., 2014). Clock genes are of particular interest in salmonids due to their association with physiological traits such as reproduction migration and smoltification (Leder et al., 2006; O’Malley et al., 2007, 2010; Paibomesai et al., 2010). Allelic diversity and variation in length polymorphism of the clock PolyQ domain was reported in four Pacific salmon species (chinook, chum, coho and pink) with overlapping geographical ranges and diversity in spawning times. This implicates clock gene duplicate may be involved in the seasonal and geographical variation in reproduction (O’Malley et al., 2007, 2010). In addition, a copy of the gene *clock* has been localised to a quantitative trait locus (QTL) responsible for 20-50% of the variation in spawning dates in female rainbow trout (Leder et al., 2006).

The circadian clock consists of intracellular transcriptional-translational feedback loops (TTFL) composed of core clock genes and stabilising accessory loop genes, which drives the rhythmic accumulation of downstream outputs, or clock-controlled genes (Partch et al., 2014; Reppert and Weaver, 2001). Circadian mechanism is highly conserved across animal species (Lowrey and Takahashi, 2011). However, deciphering the circadian clock mechanism in fish is complex. Clock and clock-related genes which are found in single copies in invertebrates such as clock, period and cryptochrome are duplicated in vertebrates (Tauber et al., 2004). In addition, salmonids were subjected to two rounds of whole-genome duplication (WGD) events (Ts3R, teleost-specific third whole-genome duplication, 320 million years ago, and Ss4R, salmonid-specific fourth whole-genome duplication, 80 million years ago) resulting in an abundance of circadian related genes (Huang, 2018; Lien et al., 2016).

The molecular mechanisms underlying circadian rhythmicity have been characterised in several model animal species including the fruit fly (*Drosophila melanogaster*), mice (*Mus musclus*), and humans (*Homo sapiens*), with relatively limited work undertaken in teleosts (Huang, 2018; Wang, 2008b) predominantly centred on zebrafish (*Danio rerio*) (Cahill, 2002; Vallone et al., 2007). The circadian system comprises all the different components by which light is perceived by the organism and is transformed into a nervous or hormonal signal (Migaud et al., 2010a). Therefore, manipulations of photic inputs and cues impact rhythms, which can be commercially exploited for aquaculture production (Migaud et al., 2010a). Research based on zebrafish has been fundamental in describing and broadly characterising clock genes and circadian rhythmicity as many findings are applicable to numerous vertebrate species. However, fish models have not yet significantly contributed to our understanding of core clock mechanisms and circadian clock control of fish physiology (Frøland Steindal and Whitmore, 2019). Teleosts represent the largest and most diverse group of vertebrates, with over 30,000 species identified to date. Each species possesses distinct characteristics and displays a considerable amount of anatomical and physiological plasticity. This is arguably the direct result of exposure to multiple, variable selection pressures caused by the highly dynamic aquatic environments they have inhabited throughout their evolutionary development (Bone, 2019). Alongside selection pressures, multiple rounds of WGD have had a large impact upon the evolution of lineages to date, with the retention of resultant gene duplicates being biased with regard to gene function (Brunet et al., 2006). Gene duplication through WGD led to redundant genes either los, non-functionalisation, with a different function than the ancestral gene (sub-functionalisation) or which acquired new functions, neo-functionalisation, (Pasquier et al., 2016), thus resulting in genome reshaping (Inoue et al., 2015). An example of this is the salmonid specific WGD (Ss4R) which preceded the origin of anadromy in salmonids, an important milestone in the evolutionary development of salmonid migration (Alexandrou et al., 2013). Whilst WGD events led to gene duplication, these duplicated genes typically resolved over time (Inoue et al., 2015). In rainbow trout, 80 to 100 million years post Ss4R, 48% of the pre-duplication ancestral genes were retained as duplicates, the remainder of the genes underwent fractionation and the duplicated protein-coding genes were lost (Berthelot et al., 2014; Lien et al., 2016). Analysis of duplicate retention in Atlantic salmon identified that 20% of duplicates from Ts3R and 55% of duplicates from Ss4R were retained as functional copies, with the prominent mechanism for duplicate loss being pseudogenisation (Lien et al., 2016). While there is a general lack of clarity surrounding the duplication and retention of functional genes post WGD, there is an unusually large complement of clock genes in salmonids (West et al., 2020).

Salmonids are amongst the most widely studied groups of fish species both scientifically and commercially, as many species of salmonid are of significant economic, societal, and environmental importance (Thorgaard et al., 2002). Many of the species within the 11 genera of the Salmonidae (Nelson et al., 2016) are of great commercial value and contribute significantly to both local and global economies through aquaculture, wild stock fisheries and recreational sport (Davidson et al., 2010; Frøland Steindal and Whitmore, 2019; Reppert and Weaver, 2001). Unlike zebrafish, salmonid species are highly seasonal in their physiology, including migration, smoltification and reproduction. Lighting and temperature manipulations are routinely used by industry to manipulate commercial broodstock ovulatory rhythm and smoltification, enabling year-round production (Migaud et al., 2013). However, understanding the intricate interactions between zeitgebers, circadian rhythmicity, seasonality, and the control of biochemical, physiological, and behavioural rhythms is complex (Migaud et al., 2010b). The completion and publication of salmonid genomes (*Salmo salar, Salvelinus alpinus, Oncorhynchus mykiss, Oncorhynchus kisutch* and *Oncorhynchus tshawytscha* to date) as part of the Functional Annotation of All Salmonid Genomes (FAASG) project (Macqueen et al., 2017), alongside RNAseq have provided great tools to study clock genes of salmonids.

The aim of this study was to identify the full complement of clock genes in Atlantic salmon (*Salmo salar*) in comparison to other commercially important salmonid species and evaluate the expression patterns of the identified genes. To do so, phylogenetic analysis of clock gene ohnologues [functional product of WGD event (Ohno, 1970)] has been performed to classify and name salmonid clock genes based on published zebrafish references and nomenclature. This was confirmed by a transcriptomic approach looking into gene expression over 24-hours in freshwater salmon kept under a controlled lighting regime. This study provides a new nomenclature for salmon clock genes that will serve as a tool for further circadian research in salmon.

## 2 MATERIALS AND METHODS

### 2.1 Identification of clock genes in salmonids with published genomes

The protein sequences of the 29 zebrafish (*D. rerio*) clock genes [*clock, arntl* (also referred to as *bmal*), *period, cryptochrome*, and *csnk1e/d* (Huang, 2018), *nr1d* (also referred to as *rev-erb*), and *ror* (Wang, 2009)] were recovered from GenBank and used as reference to interrogate the Northern pike (*Esox lucius*) [a closely related sister taxa which did not undergo the salmonid specific WGD (Macqueen et al., 2017; Rondeau et al., 2014; Varadharajan et al., 2018)] and Atlantic salmon (*S. salar*) genomes. For the benefit of this study, the core clock genes (the heterodimers forming the positive and negative feedback arms of the TTFL, *clock*:*arntl* and *period*:*cryptochrome* respectively) and accessory loop genes (individual genes *ror, nr1d*, and *csnk1* which interact with the core clock loop to either promote or repress specific heterodimer interactions) are commonly referred to collectively as clock genes. Putative core clock gene sequences were also identified for several salmonid species (*S. alpinus, O. mykiss, O. kisutch* and *O. tshawytscha*) through a combination of literature searches, BLASTp and BLASTn searches of published salmonid genomes identified as part of the Functional Annotation of All Salmonid Genomes (FAASG) initiative (Amaral and Johnston, 2012). For the benefit of this publication, they will be referred to collectively as salmonids. A BLASTp search using the default settings against the protein sequences were used for a first characterisation of the putative core clock genes. This was further refined using BLASTn using the coding sequence (CDS) against the RNA sequences (refseq rna) database and was optimised for highly similar nucleotide sequences (megablast) with an E-value below 10^−300^ and ensuring a negligible probability that the sequence was returned by chance. From the final BLASTn search, the gene and their transcriptomic isoform were aligned to the CDS of zebrafish reference genes using ClustalOmega v1.2.2 (Sievers et al., 2011).

### 2.2 Phylogenetic alignment

Amino acid sequences were aligned using GramAlign v3.0 (Russell, 2014). A Maximum Likelihood (ML) tree was inferred under the GTR model with gamma-distributed rate variation (Γ) and a proportion of invariable sites (I) using a relaxed (uncorrelated lognormal) molecular clock in RAxML (Stamatakis, 2014) with 10,000 bootstrap replicates. Gaps were handled as undetermined characters (N).

### 2.3 Classification

Nomenclature of the putative salmonid clock genes was based on phylogenetic analyses using CDS and full-length sequences to the *D. rerio* and *E. lucius* core clock genes. Genes have been renamed after the zebrafish orthologues. As a result of the salmonid specific WGD Ss4R, salmonids often have two copies of a gene which is present as a single copy in zebrafish. In most instances, this involved renaming the gene from their given predicted name. Genes were re-classified based on the nomenclature of the zebrafish reference genes (ZFIN, 2019). If a single representative was identified per species, the same name as the zebrafish reference was used. For groups with multiple representatives, the name of the orthologs were appended with numerical suffixes (.*1*, .*2*) or alphabetical suffixes (*a, b*) depending on the nomenclature of the zebrafish reference genes as a result of the previous ray fin fish WGD event, as some zebrafish genes were already denoted with alphabetical suffixes in relation to their duplication. Those with the highest percentage identity to the reference gene phylogenetically are labelled *a*. For example: the Atlantic salmon has two period 1a (per 1a) paralogs, *period 1a*.*1* (*per1a*.*1*) has the highest percentage identity when compared to the reference gene and is therefore closer to the zebrafish gene so is denoted by .*1*, and *period 1a*.*2* (*per1a*.*2*) is less identical to the ancestral form and is therefore denoted by .*2*.

### 2.4 Animal husbandry and sampling

All juvenile Atlantic salmon (*S. salar*) used in the experiment were kept at the Niall Bromage Freshwater Research Facilities at the University of Stirling. Fish (60, mean weight of 130 g, Benchmark Genetics Iceland origin) were held in an 800 L tank in a flow through system and maintained under a 12:12 Light:Darkness photoperiod (photophase from 08:00 to 20:00 using TMC AquaRay LED lamps) from 14^th^ April 2020 to 19^th^ August 2020 when sampling ended, ambient temperature ranged from 8.1 °C to 15.2 °C during this period. In the month before sampling the temperature range was 14.6 °C *±* 0.6 °C. Fish were fed daily to satiation with BioMar Orbit (2 and 3 mm pellet) during the light period using automatic feeders (Arvo-Tec, Sterner). Fish were not fed during the light period on the day of sampling. Sampling was conducted every 4-hours over a 24-hour period, starting at 10:00 on the first day and ending at 10:00 the following day (10:00[1], 14:00, 18:00, 22:00, 02:00, 06:00, 10:00[2]). At each time-point, six fish were sampled (Supplementary Table S1). Following lethal anaesthesia (MS222), brains were dissected out and snap frozen directly in liquid nitrogen and stored at −80 °C until analysis.

### 2.5 RNA extraction and sequencing

RNA was isolated from the whole brain using TRI reagent (Sigma, St Louis, MO, USA) and RNA concentration was tested using a Qubit 2.0 Fluorometer (ThermoFisher Scientific, Waltham, MA, USA). The 42 RNA samples were submitted to Novogene UK (Cambridge) for RNA sequencing. Samples were submitted to quality control (Illumina BioAnalyzer®) revealing the RNA integrity number (RIN) of samples valued between 8.8-10.0. For each sample (900 ng), a library was prepared using NEB Next^®^ UltraTM RNA Library Prep Kit and processed and sequenced using Illumina NovaSeq^®^ 6000 S4 PE150 (6 GB of data per sample, ca. 40 million reads).

### 2.6 RNA sequencing analysis

Clean reads were obtained from the raw reads by filtering ambiguous bases, PCR duplicates, low quality sequences (*<* Q20), length (150 nt), absence of primers/adaptors and complexity (entropy over 15) using fastp (Chen et al., 2018). Ribosomal RNA was further removed using SortMeRNA v3.0.2 (Kopylova et al., 2012) against the Silva version 119 rRNA databases Quast et al. (2012). The remaining reads were aligned to the annotated *S. salar* genome ICSASG v2.99 (Accession GCA 000233375.4) using HiSat2 v2.2.0 (Kim et al., 2019). The expression levels were estimated based on the genome annotation using StringTie2 v2.1.0 (Kovaka et al., 2019) following the workflow: (a) for each sample, map the reads to the genome with HiSat2 and assemble the read alignments with StringTie2; (b) merge the assemblies in order to generate a non-redundant set of transcripts observed in all the samples; (c) for each sample, estimate transcript abundances and generate read coverage tables expressed in the fragments per kilobase of exon per million mapped reads (FRKM).

### 2.7 Statistical analysis

All tests and analysis were performed using R v4.1 (R Core Team, 2020). The expression values were scaled in Transcripts Per Millions (TPM) before been normalised by size factor using the median ratio method described by Anders and Huber (2010) using DESeq2 v1.32.0 (Love et al., 2014). Differential expression was estimated using the function *lfcShrink* (Stephens, 2017). The heatmaps were created from the normalised count files and scaled by row to highlight individual gene expression. Daily rhythms across the genome including identified clock genes were identified from the 24-hour dataset using RAIN v1.26.0 (Thaben and Westermark, 2014) and MetaCycle v1.2.0 (Wu et al., 2016) implementation of JTKCycle (Hughes et al., 2010) and a threshold *p*-value of 0.001 and a minimum relative oscillation amplitude of 10%. All p-value reported were corrected for False Discovery Rate (FDR) using Bonferroni adjustment.

## 3 RESULTS

### 3.1 Identification of clock genes in salmonids with published genomes

From the BLASTn, 143 clock gene variants were returned; a substantial number were highly similar predicted variants of the same gene and had identical EMBL accession numbers (LOC ID) and the CDS were over 90% identity. Most of the differences were found in the UTR. The CDS of variants sharing LOC ID were aligned to the CDS of *D. rerio* reference genes. Variants with the highest percentage identity compared to the reference for each locus were selected, leaving a total of 48 core clock genes identified in the *S. salar* across the 7 gene families explored (Table 1 and Supplementary Table S2).

**Table 1.** Genomic structure of the gene families associated with the circadian clock of salmonids. Phylogenetic trees upon which the table was based on can be found in Figure 1 and Supplementary Figures S1-S7. Key: • gene detected in genome; *\** gene not detected in genome; gene not detected in genome but identify from transcriptomes data. DR *Danio rerio* (GCA 000002035.4), EL *Esox lucius* (GCA 011004845.1), SSA *Salmo salar* (GCA 000233375.4), SA *Salvelinus alpinus* (GCA 002910315.2), OM *Oncorhynchus mykiss* (GCA 002163495.1), OK *Oncorhynchus kisutch* (GCA 002021735.2), OT *Oncorhynchus tshawytscha* (GCA 002872995.1).

| Family | Genes | DR | EL | SSA | SA | OM | OK | OT |
| --- | --- | --- | --- | --- | --- | --- | --- | --- |
| *clock* | *clock1a* | ● | ● | ●● | ●● | ●● | ●● | ●● |
|  | *clock1b* | ● | ● | - | - | - | - | - |
|  | *clock2 (npas2)* | ● | ● | ●● | ●● | ●● | ●● | ● |
| *arntl* | *arntl1a* | ● | ● | ●● | ●● | ●● | ●● | ●● |
|  | *arntl1b* | ● | ● | ●● | ● | ●● | ●● | ● |
|  | *arntl2* | ● | ●● | ●●● | ●● | ●●● | ●●● | ●● |
| *period* | *per1a* | ● | ● | ●● | ●● | ●● | ●● | ●● |
|  | *per1b* | ● | ● | ● | ● | ● | ● | ● |
|  | *per2* | ● | ● | ●● | ●● | ●● | ●● | ●● |
|  | *per3* | ● | ● | ● | ● | - | - | - |
| *cryptochrome* | *cry1a* | ● | ● | ●● | ●● | ●● | ●● | ●● |
|  | *cry2* | ● | ● | ● | ● | ● | ● | * |
|  | *cry3a* | ● | ● | - | - | - | - | - |
|  | *cry3b* | ● | ● | ●● | ●● | ● | ● | * |
|  | *cry4* | ● | ● | ● | - | - | - | - |
|  | *cry5* | ● | ● | ● | ● | ● | ● | ● |
| *nr1d* | *nr1d1* | ● | ● | ●● | ●● | ●● | ●● | ●● |
|  | *nr1d2a* | ● | ● | ●● | ●● | ●● | ● | ●● |
|  | *nr1d2b* | ● | ● | ●● | ●● | ●● | ●● | ●● |
|  | *nr1d4a* | ● | ● | ●● | ●● | ●● | ●● | ●● |
|  | *nr1d4b* | ● | ● | ●● | ●● | ●● | ●● | ●● |
| *ror* | *roraa* | ●● | ●● | ●● | ●● | ●● | ●● | ●● |
|  | *rorab* | ● | - | - | - | - | - | - |
|  | *rorb* | ● | ● | ●● | ●● | ●● | ●● | ●● |
|  | *rorca* | ● | ● | ●● | ●● | ●● | ●● | ●● |
|  | *rorcb* | ● | ● | ●● | ●● | ●● | ● | ●● |
| *csnk1e/d* | *csnk1e* | ● | ● | ●●●● | ●●●● | ●●●● | ●● | ●●●● |
|  | *csnk1da* | ● | ● | - | - | - | - | - |
|  | *csnk1db* | ● | ● | ●● | ● | ●● | ●● | ●● |

### 3.2 Interpretation and classification

All the core clock and clock accessory loop genes were identified and classified (Table 1). Gene members of each family were aligned together in a gene-family approach to aid clarification of paralog nomenclature.

The *period* (*per*) family is used as a typical example (Figure 1), showing the resulting gene tree from the identification and classification of per genes in salmonids indicating reciprocal gene retention (red), gene duplication (orange), and reciprocal gene loss (blue).

**Figure 1.**
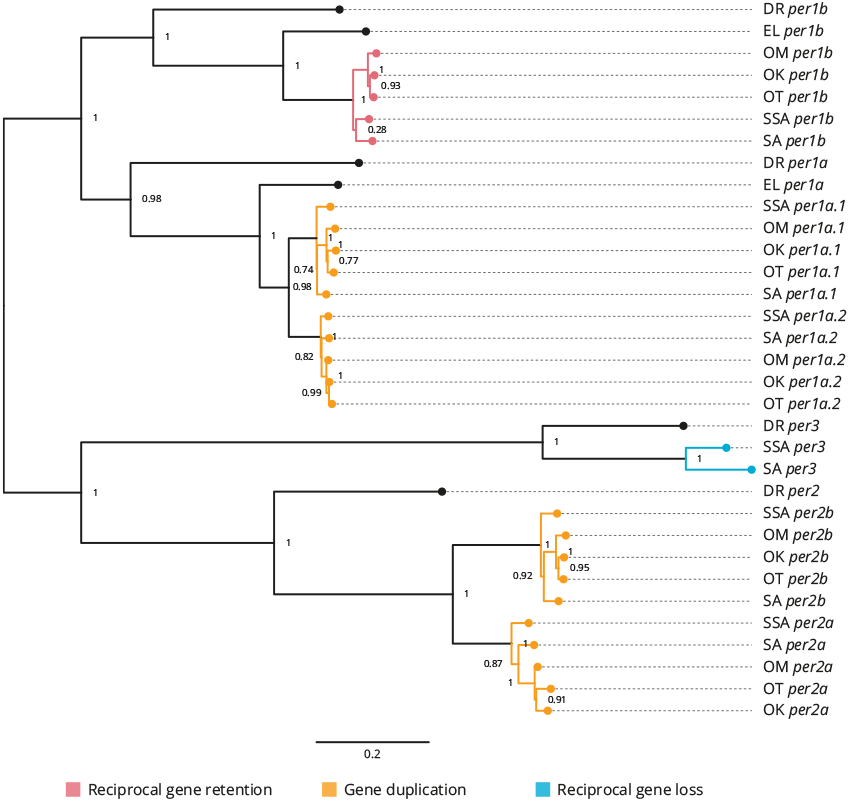
Phylogenetic relationship of the period clock gene family (*per*). Gene family members from *Danio rerio* (DR), *Esox Lucius* (EL), *Salmo salar* (SSA), *Salvelinus alpinus* (SA), *Oncorhynchus mykiss* (OM), *Oncorhynchus kisutch* (OK) and *Oncorhynchus tshawytscha* (OT) were re-annotated. Values on the relevant node depict the bootstrap values. Reciprocal gene retention (red), gene duplication (orange) and reciprocal gene loss (blue).

#### 3.2.1 Period (Figure 1 and Supplementary Figure S1)

*per1a* is duplicated across all salmonids (*per1a.1, per1a.2*). *per1b* is retained in all salmonids (*per1b.1, per1b.2*). *per2* appears to be duplicated after the latest common ancestor *E. lucius* (*per2a, per2b*). *per3* is retained in *S. salar* and *S. alpinus*, but appears to have been lost in *O. mykiss, O. kisutch*, and *O. tshawytsch*a.

#### 3.2.2 Clock (Supplementary Figure S2)

*clock1a* is duplicated in all salmonids (*clock1a.1, clock1a.2*). *clock1b* was lost compared to *D. rerio* and *E. lucius* and is not present in the salmonids. *clock2* (*npas2*) is duplicated across all salmonids analysed (*clock2a, clock2b*) with a differential ohnolog loss in *O. tshawytscha* (aligns to *clock2a*).

#### 3.2.3 Aryl hydrocarbon receptor nuclear translocator-like (Supplementary Figure S3)

*arntl1a* is duplicated in all salmonids (*arntl1a.1, arntl1a.2). arntl1b* is duplicated in all salmonids (*arntl1b.1, arntl1b.2*) with differential ohnolog loss in both *S. alpinus* (aligns to *arntl1b.2*) and *O. tshawytscha* (algins to *arntl1b.1*). There appears to be a duplication of *arntl2* in the latest common ancestor which is retained across all salmonids (*arntl2a, arntl2b*). There is a third copy identified in *S. salar, S. alpinus, O. mykiss* and *O. kisutch* (*arntl2c*).

#### 3.2.4 Cryptochrome (Supplementary Figure S4)

*cry1a* is duplicated across all salmonids (*cry1a.1, cry1a.2*). *cry1b* appears to be lost after *D. rerio. cry2* is retained in all salmonids with the exception of *O. tshawytscha* as there is currently not enough information available regarding *cry2* in the species. There is an apparent gene loss of *cry3a* in salmonids after the most recent common ancestor, *E. lucius. cry3b* is duplicated in *S. salar* and *S. alpinus* (*cry3b.1, cry3b.2*) there is an apparent gene loss in *O. mykiss* and *O. kisutch* (aligns to *cry3b.1*), currently there is not enough information available regarding *cry3b* in *O. tshawtscha*. There is an apparent gene loss of *cry4* in all salmonids after the latest common ancestor *E. Lucius* in all species of salmonid except from *S. salar. cry5* is retained in all salmonids.

#### 3.2.5 Nuclear receptor subfamily 1 group d (Supplementary Figure S5)

*nr1d1* is duplicated across all salmonids (*nr1d1a, nr1d1b*). *nr1d2a* is duplicated across all salmonids (*nr1d2a.1, nr1d2a.2*), with an apparent differential ohnolog loss in *O. kistuch* (aligns to *nr1d2a.2*). *nr1d2b* is duplicated across all salmonids (*nr1d2b.1, nr1d2b.2*). *nr1d4a* is duplicated across all salmonids (*nr1d4a.1, nr1d4a.2*). *nr1d4b* is duplicated across all salmonids (*nr1d4b.1, nr1d4b.2*).

#### 3.2.6 RAR-related orphan receptor (Supplementary Figure S6)

*roraa* is duplicated across all salmonids (*roraa*.1, *roraa.2*), there is an apparent differential gene loss in *O. tshawytscha* (aligns to *roraa.1*) and an apparent loss of both ohnologs in *O. kisutch*. There is an apparent loss of *rorab* after the latest common ancestor, *E. lucius. rorb* has been duplicated across all salmonids (*rorb.1, rorb.2*). *rorca* has been duplicated across all salmonids (*rorca.1, rorca.2*). *rorcb* has been duplicated across all salmonids (*rorcb.1, rorcb.2*) with an apparent differential ohnolog loss in *O. kisutch* (aligns to *rorcb.2*).

#### 3.2.7 Casein kinase 1 delta (Supplementary Figure S7)

*csnk1db* appears to be duplicated across all salmonids (*csnk1db.1, csnk1db.2*) apart from *S. alpinus* which displays an apparent differential ohnologs loss (aligns to *csnk1db.1*).

#### 3.2.8 Casein kinase 1 epsilon (Supplementary Figure S7)

There appears to be multiple duplications of csnk1e across all the salmonids, resulting in four csnk1e paralogs (*csnk1ea, csnk1eb, csnk1ec, csnk1ed*). With *O. kisutch* displaying a potential differential ohnologs loss (aligns to *csnk1ea, csnk1eb*).

### 3.3 Transcriptomic Analysis

In total, 986,545,058 raw reads were sequenced for 42 samples (Supplementary Table S2). The reads were deposited in the European Bioinformatics Institute (EBI) European Nucleotide Archive (ENA) project ID PRJEB41327. After filtering, 975,284,560 clean reads (98.86%) passed the mRNA cleaning step and were used for the following process. Of the clean reads, 97.71% were aligned to the published *S. salar* genome ICSASG v2.99 (Accession GCA 000233375.4). A total of 55,819 distinct genes were recovered (Supplementary Data S1). All clock genes classified in the *in silico* gene identification were recovered.

### 3.4 Rhythmic and circadian gene expression

Significantly rhythmically expressed genes were identified using RAIN and MetaCycle with JTK analysis (Supplementary Data S2). Various thresholds were evaluated (Supplementary Table S3). From the 48 genes clock genes, 16 were significantly rhythmically expressed (RAIN analysis, *p <* 0.001, relative amplitude *≥* 10%), of which 11 also had a significant circadian expression patten over a 24-hour period (JTK analysis, *p <* 0.001, relative amplitude *≥* 10%, Figure 2A and Supplementary Figures S8 & S9). Details of the gene family are plotted in Figure reffig:figure2B. From the six period genes identified five were highly significantly rhythmically expressed (*per1a.1, per1a.2, per1b, per2a, per2b*), including three that also exhibited circadian expression pattern (*per1a.1, per1a.2, per2b*). The acrophase for *per1a.1* and *per1a.2* are in phase at 06:00, *per1b* has a negative phase shift in comparison to *per1a* paralogs at 08:30. *per2a* and *per2b* are out of phase by two hours, with *per2a* peaking at 03:00 and *per2b* at 01:00.

**Figure 2.**
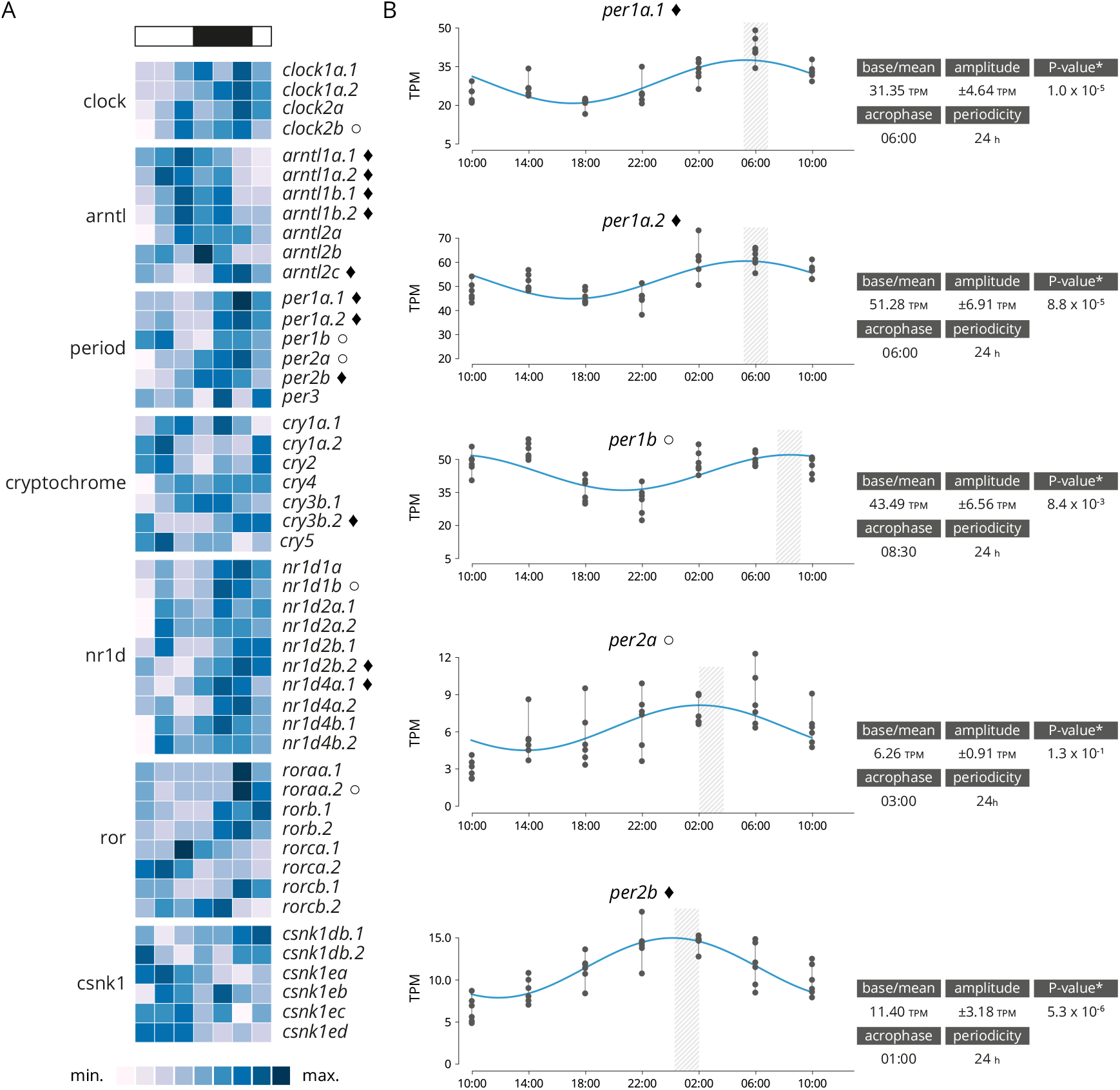
**(A)** Heatmap displaying average diel expression of identified clock genes under constant LD (12:12, n = 6 per time point). The heatmap of the relative expression of each individual gene [scaled from lowest expression to highest expression]. Black diamond indicates significantly cyclic gene (*p <* 0.001) [JTK and RAIN analysis], White circles denote rhythmic genes (*p <* 0.001) [RAIN analysis]. **(B)** Significantly clock gene expression. Parameters of the cyclic sin-cosine function calculated by MetaCycle with JTK for diel expression of clock genes in brains collected from Atlantic salmon smolt exposed to an LD 12:12 photoperiod.

Overall, 2,864 genes exhibited a rhythmic expression pattern (rhythmic peak; RAIN analysis, *p <* 0.001; Supplementary Data S2), and of which 1,215 genes showed a significance circadian expression patten (peak and trough; JTK analysis; *p <* 0.001). Rhythmic gene were clustered by acrophase (expression peak) synchronicity (Figure 3). The number of circadian expressed genes with the same acrophase range oscillate between 95 and 419. The majority of genes are expressed in one of two peaks at 10:00 [first sampling in the light] and 22:00 [first sampling in the darkness]. The highly significantly rhythmically expressed clock genes are distributed similarly to the rest of those which are significantly expressed in a circadian pattern (p = 0.15). The majority are peaking at 22:00. Sub-figures exclude 3 highly expressed genes that bias the scale of the graphic: Ependymin-1 (*>* 10,000 TPM) Ependymin-2 (*>* 5,000 TPM) and calcium voltage-gated channel auxiliary subunit alpha 2 delta 4 (*>* 2,000 TPM); all three genes are related with central nervous system plasticity and memory formation.

**Figure 3.**
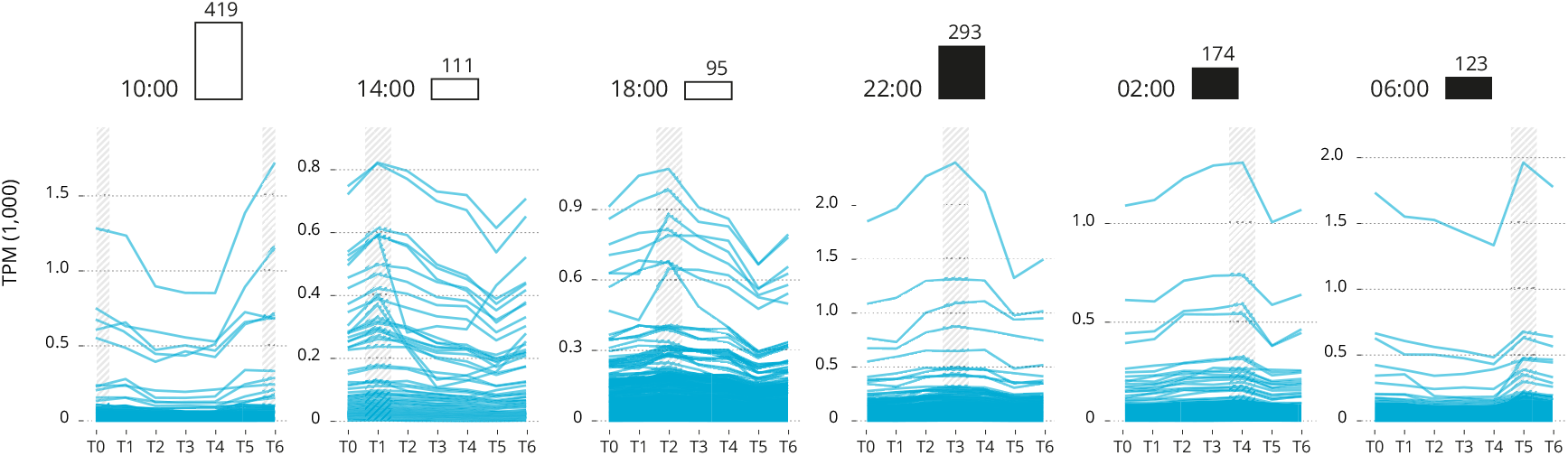
Genes groups based on their acrophase (expression peak) synchronicity. The number of gene is reported over the barplot (white and black boxes denote light or darkness conditions respectively). Sub-figures exclude three highly expressed genes that bias the scale of the graphic: Ependymin-1 (*>* 10,000 TPM) Ependymin-2 (*>* 20,000 TPM) and calcium voltage-gated channel auxiliary subunit alpha 2 delta 4 (*>* 2,000 TPM).

## 4 DISCUSSION

In the present study, we have identified and characterised 48 core clock and accessory loop genes *in silico* in several salmonid species with published genomes using the latest common ancestors as reference points. For each of the salmonid species with published genomes, the core clock and accessory loop genes were identified and characterised *in silico*. There is a differential retention of genes originating in the latest common ancestors, *E. lucius* and *D. rerio*. In addition to this, there are numerous ohnologs (Ohno, 1970) which are consistent with the salmonid specific whole genome duplication event Ss4R (Lien et al., 2016). There is also evidence of differential gene loss in salmonids based on the current gene annotations (Table 1). Findings from this study further highlight the rich complexity of core clock genes previously outlined in *S. salar* (West et al., 2020), and is displayed across a wider complement of salmonid species. The gene families identified differ slightly to those explored by West et al.(2020) as the focus of this study was to identify all members of gene families *clock, arntl, period, cryptochrome, ror, nr1d* and *csnk1*. Identified genes common to both studies coincide with one another and paralog pairs identified using ML phylogenetic alignments are also identified as paralogs (West et al., 2020).

In comparison to *D. rerio* with 29 reference clock genes (Wang, 2008b; Huang et al., 2010; Toloza-Villalobos et al., 2015), most clock genes identified in salmonids appear to have been duplicated. In addition to these lineage-specific duplications there is also evidence of differential ohnolog loss in some species of salmonids based on the current genome annotations, *per1b, cry2*, and *cry5* appear to have been retained across all species from the latest common ancestors, *per3* appears to only be retained in *S. salar* and *S. alpinus* and *cry4* appears to only be retained in *S. salar*. Whereas *clock1b, cry1b, cry3a, csnk1da* all appear to have been lost in salmonids after the latest common ancestor *E. lucius*. Some species of salmonids have better genome annotations than others, and currently it is not possible to identify the effects of the WGD for every core clock gene family in *O. tshawytscha* at present. As expected across most of the gene families except where there is limited information, the *Oncorhynchus* spp. tend to appear together on the same clade with *S. salar* and *S. alpinus* commonly grouped together in a sister clade. A typical example of the clock gene families identified is the cryptochrome family. *cry1a* is one of the major clades in the cryptochrome family, as a result of duplication there are three main paralogous subfamilies *cry1a, cry1b* and *cry1c. cry1a* and *cry1b* are phylogenetically related to tetrapods, with *cry1a* being the most conserved of the two subfamilies (Mei et al., 2015). Zebrafish possess both cry1a and *cry1b* however, all the salmonid species only appear to have inherited *cry1a* which has been duplicated as a result of the WGD, resulting in two paralogs *cry1a.1* and *cry1a.2*. Except for *O. tshawytscha* (for which there is not currently enough information in the genome annotation) *cry3b* is duplicated in all salmonid species with an apparent gene loss in both *O. kisutch* and *O. mykiss*. This is suggestive of the non-functualisation and subsequent loss of *cry3b.2* in the *Oncorhynchus* spp. As a result of the Ss4R WGD clock most genes appear to be duplicated in salmonids with some exceptions that appear to have been lost in several of the salmonids. As annotations improve, it is thought that additional gene losses will become more apparent. This study gives insight into the post duplication effects on clock gene family members and provides a fundamental tool for further circadian work in salmonids.

Clock genes interact with each other, generating oscillations in gene expression. Their underlying principle is to create successive gene activation in the form of a cycle, forming an autoregulatory feedback loop which perpetually cycles approximately every 24 hours (Ripperger and Albrecht, 2009). This in turn influences downstream targets, whose time-of-day specific expression is determined by the central circadian mechanism (Ripperger and Albrecht, 2009). This study confirmed the expression of the identified clock genes in the brain of *S. salar* smolts over a 24-hour period and showed that 11 out of the 48 core clock genes were highly significantly (*p <* 0.001) rhythmically expressed. We used a two complementary approach, RAIN which allowed us to detect accurately rhythms in time series irrespectively of the shape of the expression pattern, and JTK with a focus on circadian expression patterns. The vast majority of the JTK findings are included in the RAIN results (Supplementary Table S3). This allowed us to distinguish between rhythmic expressed genes and circadian pattern expressed genes.

The period family is particularly interesting, as except for one gene (*per3*) all the genes were significantly rhythmically expressed. A difference in acrophase between paralogs was also identified. There was a clear difference in expression pattern observed between *per1a* paralogs (*per1a.1, per1a.2*) and *per1b*, with *per1a* peaking at 06:00 displaying a positive phase shift in comparison to *per1b* which peaks at 08:30. This positive phase shift coincides with results observed in zebrafish, it was reported that *per1a* and *per1b* paralogs displayed a shifted phase of gene expression in zebrafish with *per1a* peaking 4 hours prior to *per1b* under a 12:12 LD lighting regime (Amaral and Johnston, 2012). In zebrafish, *per1* paralogs were shown to display a distinct difference in spatial and temporal expression in the brain, providing strong evidence for the duplicate pair to have undergone sub- or neo-functionalisation (Wang, 2008a). The duplicated Atlantic salmon *per1a* paralogs (*per1a.1* and *per1a.2*) share the same periodicity and acrophase, thus indicating that both genes may share a similar functionality. This indicates that the paralogs may have undergone sub-functionalisation - in which the ancestral functions have become partitioned and each paralog potentially has particular adaptations for different tissues, developmental stages or environmental conditions (Innan, 2009). In mammalian literature, it is widely reported that *Cry1/2* form heterodimers with *Per1/2/3* (Rosensweig et al., 2018). *However, in this study, the only highly significantly rhythmically entrained cryptochrome* genes was the *cry3b.2* paralog which peaked two hours before the closest *period* family member (*per2b*) at 20:30, in agreement with the pattern of *cry2* expression identified in zebrafish 12-hours after light onset (Hirayama et al., 2019). In mammals, *Per2* is said to repress *Nr1d2* transcription thus upregulating *arntl1* transcription (Chiou et al., 2016), in keeping with these findings in this study *nr1d2b.2* peaks at 08:30, around 10 hours before peak expression of *arntl1a.1/1a.2/1b.1* (18:00) and *per2b* peaks at 01:00. This suggests that the relationship between clock gene paralogs in salmonids may be similar to that of those in mammalian species.

How salmonids fit into the widely accepted mechanism for circadian rhythmicity is difficult to evaluate due to the limited understanding surrounding circadian mechanisms in fish in general and the complexity are yet to be exploited to the full potential (Frøland Steindal et al., 2018), there are still many unknowns surrounding the mechanism in salmonids as it has been far less explored than other species of teleost. In comparison to species such as *D. melanogaster, M. musclus, H. sapiens* and even *D. rerio* there has been limited work undertaken in salmonids to further identify clock genes and their respective roles within the circadian system (Frøland Steindal et al., 2018). So far, the expression of individual clock genes has been previously investigated in salmonids (Davie et al., 2009; Huang et al., 2010; West et al., 2020). Clock gene member identification has previously been hampered by paralogs with high sequence similarity, which does not easily allow for individual identification by qPCR. Individual gene identification using RNA sequencing has allowed for paralogs to be better classified and individual gene expression patterns ascertained. Although, single gene duplication events in mammalian species have enabled the evolution of specialisation and regulatory sophistication in the temporal regulation of local physiology (Looby and Loudon, 2005); the core clock genes associated with circadian rhythmicity remains largely conserved across a diverse range of organisms spanning vast evolutionary time periods (Bell-Pedersen et al., 2005; Cox and Takahashi, 2019). This study supports that whilst the compliment of clock genes is far richer in salmonids, the function of core clock genes remains conserved and are therefore likely function similarly to other more studied organisms. Some salmonid species display reciprocal gene losses post Ss4R, but many of clock genes identified appear to be duplicated in the salmonid species investigated. The existence of such paralogs whilst increasing genetic complexity may enable gene family members to specialise and extend their ancestral role, which can lead to a shift in the identity of components of the molecular clock (Looby and Loudon, 2005; West et al., 2020). Ambient temperature affects gene expression and physiology in ectotherms; in zebrafish, the clock seems to be temperature-compensated, changing the amplitude of some critical clock genes (Lahiri et al., 2005). However, the influence of temperature on the clock system in salmonids remains poorly studied and should be investigated further.

Whilst this study has furthered the identification of clock gene family members in central salmonids, additional work is required to further elucidate the complexity of the circadian mechanism and how the complement of clock genes identified individually function as components of this mechanism. It is important to note that alongside individual variability, the whole brain being analysed will have influenced gene expression levels and rhythmicity due to the highly decentralised organisation of clock genes in teleosts and localisation of clock gene expression in specific regions of the brain (Moore and Whitmore, 2014). This study provides a fundamental tool to explore the role of the enriched number of clock genes related to seasonal driven life history transition in salmonids.

## DATA AVAILABILITY STATEMENT

The raw sequencing reads of all libraries are available from EBI/ENA via the project PRJEB41327.

## ETHICS STATEMENT

Animals were treated in accordance with the UK Animals (Scientific Procedures) Act 1986 Amended Regulations (SI 2012/3039) and the experiment was approved by the Animal Welfare and Ethical Review Body of the University of Stirling (AWERB/19 20/097/).

## AUTHOR CONTRIBUTIONS

CMB, JVH, MB and HM: study conception and design. CMB and HM: sample and data acquisition. CMB, MB and ME: data analysis. HM and JVH: funding acquisition. CMB, HM, ME and MB drafting and editing of the manuscript. All authors contributed to the article and approved the submitted version.

## ACKNOWLEDGMENTS

Thank you to all reviewers for their constructive comments. We also thank staff at the Niall Bromage Freshwater Research Facilities for looking after the experimental fish.

