## Supplementary Figures S1-S9 and Supplementary Tables S1-S9 for "Rhythmic clock gene expression in Atlantic salmon parr brain"

#### ABSTRACT

To better understand the complexity of clock genes in salmonids, a taxon with an additional whole genome duplication, an analysis was performed to identify and classify gene family members (*clock*, *arntl*, *period*, *cryptochrome*, *nr1d*, *ror*, and *csnk1*). The majority of clock genes, in zebrafish and Northern pike, appeared to be duplicated. In comparison to the 29 clock genes described in zebrafish, 48 clock genes were discovered in salmonid species. There was also evidence of species-specific reciprocal gene losses conserved to the *Oncorhynchus* sister clade. From the six period genes identified three were highly significantly rhythmic, and circadian in their expression patterns (*per1a.1*, *per1a.2*, *per1b*) and two was significantly rhythmically expressed (*per2a*, *per2b*). The transcriptomic study of juvenile Atlantic salmon (parr) brain tissues confirmed gene identification and revealed that there were 2,864 rhythmically expressed genes ($p < 0.001$), including 1,215 genes with a circadian expression pattern, of which 11 were clock genes. The majority of circadian expressed genes peaked two hours before and after daylight. These findings provide a foundation for further research into the function of clock genes circadian rhythmicity and the role of an enriched number of clock genes relating to seasonal driven life history in salmonids.

#### SUPPLEMENTARY MATERIAL

##### Data S1

The expression level. For each sample, estimate gene abundance expressed in the fragments per kilobase of exon per million mapped reads (FRKM) as exported by StringTie2/HiSat2. (CSV)

##### Data S2

All rhythmically expressed genes (after Transcripts Per Million (TPM) scaling and normalisation). (CSV)

#### SUPPLEMENTARY FIGURES

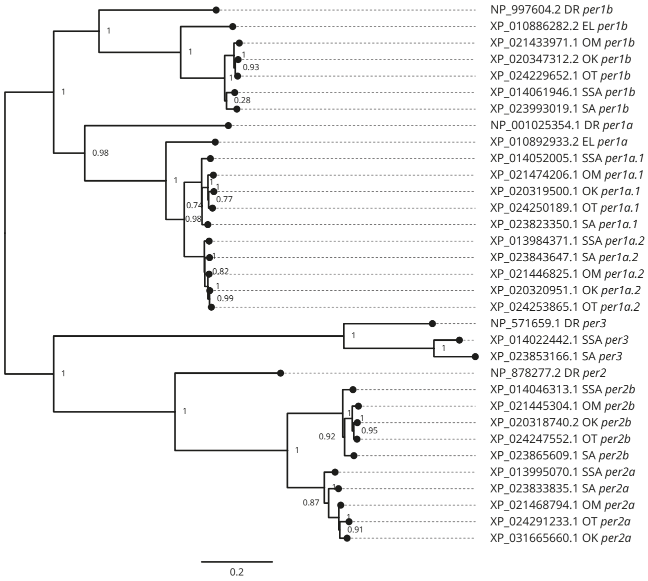


**Figure S1.** Phylogenetic relationship of the *period* gene family. Values on the relevant node depict the bootstrap values. Sequence ID used and identified are provided.

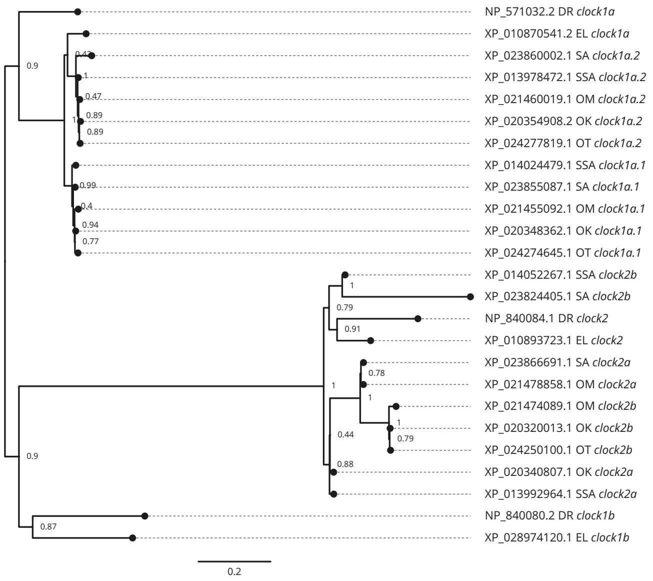


**Figure S2.** Phylogenetic relationship of the *clock* gene family. Values on the relevant node depict the bootstrap values. Sequence ID used and identified are provided.

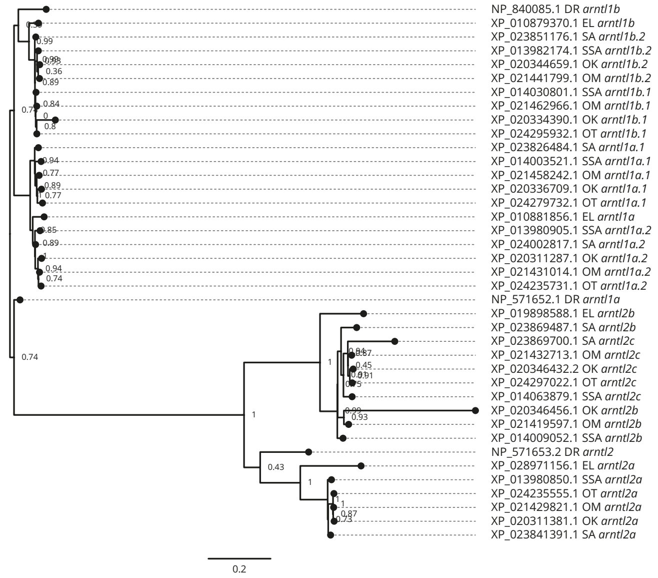


**Figure S3.** Phylogenetic relationship of the *aryl hydrocarbon receptor nuclear translocator-like* gene family (*arntl*). Values on the relevant node depict the bootstrap values. Sequence ID used and identified are provided.

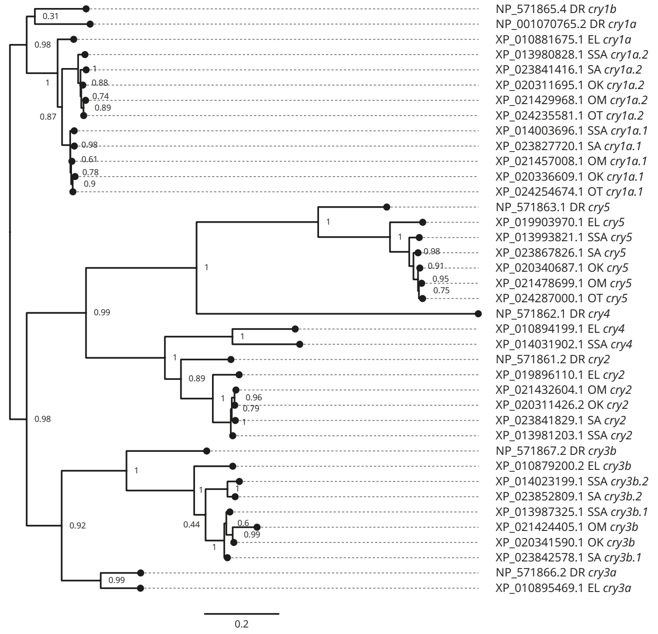


**Figure S4.** Phylogenetic relationship of the *cryptochrome* gene family (*cry*). Values on the relevant node depict the bootstrap values. Sequence ID used and identified are provided.

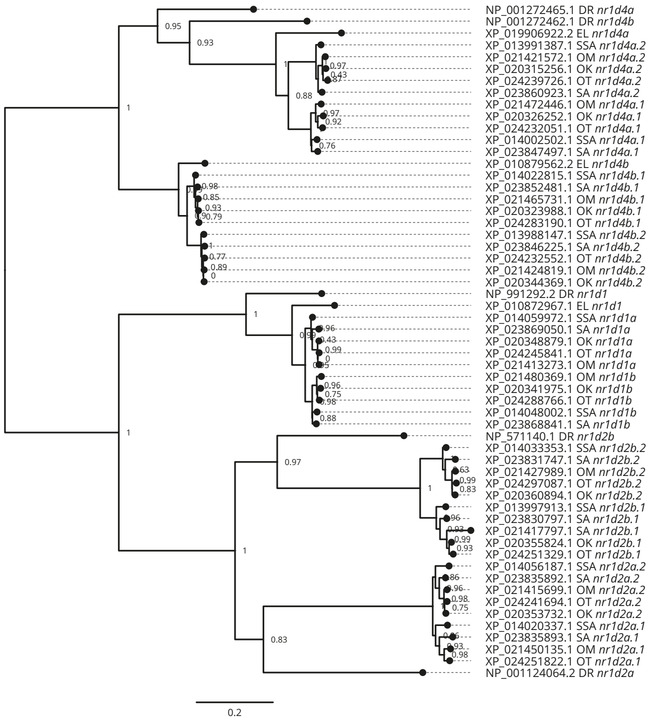


**Figure S5.** Phylogenetic relationship of the *nuclear receptor subfamily 1 group d* gene family (*nr1d*). Values on the relevant node depict the bootstrap values. Sequence ID used and identified are provided.

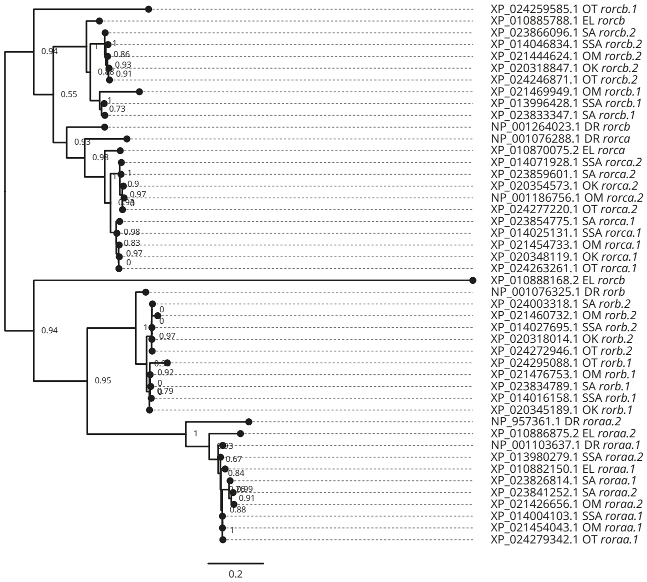


**Figure S6.** Phylogenetic relationship of the *RAR-reticulated orphan receptor* (*ror*). Values on the relevant node depict the bootstrap values. Sequence ID used and identified are provided.

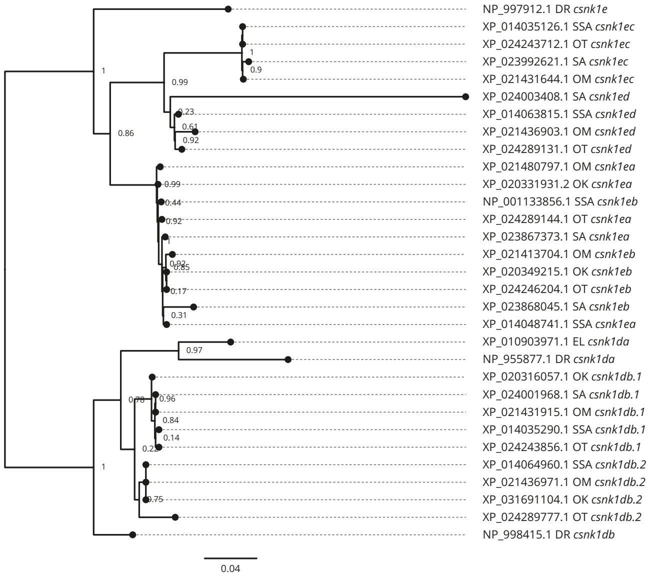


**Figure S7.** Phylogenetic relationship of the *casein kinase 1 delta* (*csnk1d*) and *epsilon* (*csnk1e*). Values on the relevant node depict the bootstrap values. Sequence ID used and identified are provided.

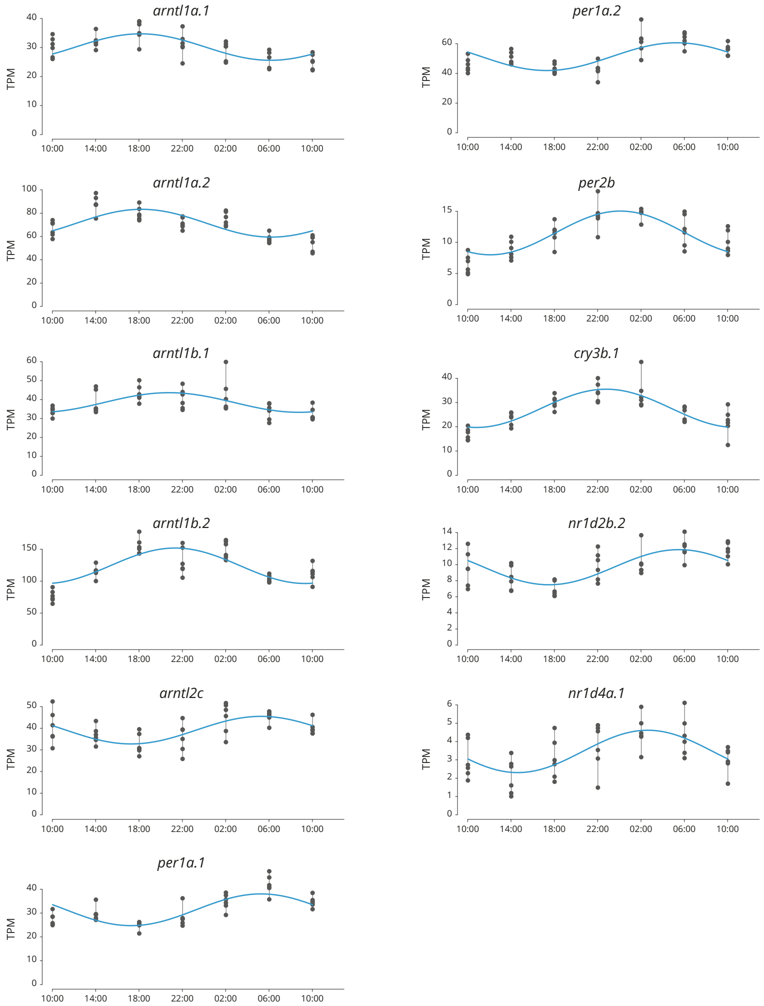


**Figure S8.** Significantly cyclical gene expression. Parameters of the cyclic sin-cosine function calculated by MetaCycle with JTK ($P < 0.001$) for diel expression of clock genes in brains collected from Atlantic salmon smolt exposed to an LD 12:12 photoperiod.

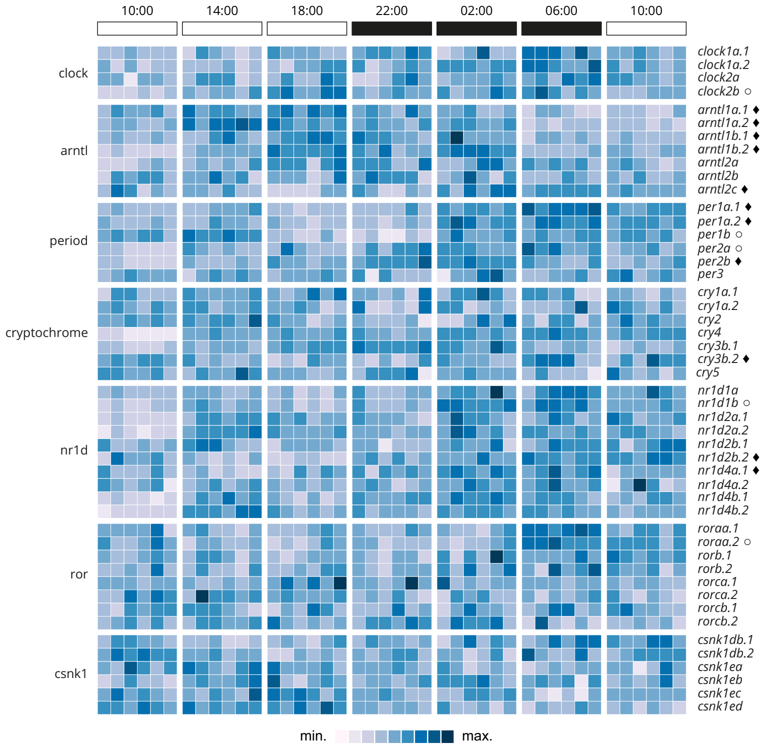


**Figure S9.** Heatmap displaying individual diel expression of identified clock genes under constant LD (12:12, n = 6 per time point). The heatmap of the relative expression of each individual gene [scaled from lowest expression to highest expression]. Black diamond indicates significantly cyclic gene ($p < 0.001$) [JTK and RAIN analysis], White circles denote rhythmic genes ($p < 0.001$) [RAIN analysis].

### SUPPLEMENTARY TABLES

**Table S1.** Samples and reads details.

| Sample | SampleID | BrainID | Sample EBI Accession | Raw Reads | Date | Time | Timepoint | Mapping |
|---|---|---|---|---|---|---|---|---|
| A1 | 2/LD/T0/1 | 85 | ERS5329244 | 47723702 | 2020-08-24T09:00:00Z | 0 | T0 | 97.8% |
| A2 | 2/LD/T0/2 | 86 | ERS5329245 | 41776194 | 2020-08-24T09:00:00Z | 0 | T0 | 98.0% |
| A3 | 2/LD/T0/3 | 87 | ERS5329246 | 48551618 | 2020-08-24T09:00:00Z | 0 | T0 | 97.9% |
| A4 | 2/LD/T0/4 | 88 | ERS5329247 | 40458840 | 2020-08-24T09:00:00Z | 0 | T0 | 98.0% |
| A5 | 2/LD/T0/5 | 89 | ERS5329248 | 39687614 | 2020-08-24T09:00:00Z | 0 | T0 | 97.5% |
| A6 | 2/LD/T0/6 | 90 | ERS5329249 | 47388666 | 2020-08-24T09:00:00Z | 0 | T0 | 97.9% |
| A7 | 2/LD/T1/1 | 97 | ERS5329250 | 48544056 | 2020-08-24T13:00:00Z | 4 | T1 | 97.9% |
| A8 | 2/LD/T1/2 | 98 | ERS5329251 | 42958682 | 2020-08-24T13:00:00Z | 4 | T1 | 98.0% |
| A9 | 2/LD/T1/3 | 99 | ERS5329252 | 48892118 | 2020-08-24T13:00:00Z | 4 | T1 | 97.9% |
| A10 | 2/LD/T1/4 | 100 | ERS5329253 | 57346024 | 2020-08-24T13:00:00Z | 4 | T1 | 98.0% |
| A11 | 2/LD/T1/5 | 101 | ERS5329254 | 44973308 | 2020-08-24T13:00:00Z | 4 | T1 | 97.9% |
| A12 | 2/LD/T1/6 | 102 | ERS5329255 | 46567214 | 2020-08-24T13:00:00Z | 4 | T1 | 97.8% |
| A13 | 2/LD/T2/1 | 109 | ERS5329256 | 46177414 | 2020-08-24T17:00:00Z | 8 | T2 | 97.9% |
| A14 | 2/LD/T2/2 | 110 | ERS5329257 | 49718214 | 2020-08-24T17:00:00Z | 8 | T2 | 97.8% |
| A15 | 2/LD/T2/3 | 111 | ERS5329258 | 40568918 | 2020-08-24T17:00:00Z | 8 | T2 | 97.7% |
| A16 | 2/LD/T2/4 | 112 | ERS5329259 | 47615560 | 2020-08-24T17:00:00Z | 8 | T2 | 97.8% |
| A17 | 2/LD/T2/5 | 113 | ERS5329260 | 49860608 | 2020-08-24T17:00:00Z | 8 | T2 | 97.8% |
| A18 | 2/LD/T2/6 | 114 | ERS5329261 | 45925090 | 2020-08-24T17:00:00Z | 8 | T2 | 97.4% |
| A19 | 2/LD/T3/1 | 121 | ERS5329262 | 49124232 | 2020-08-24T21:00:00Z | 12 | T3 | 97.2% |
| A20 | 2/LD/T3/2 | 122 | ERS5329263 | 46414190 | 2020-08-24T21:00:00Z | 12 | T3 | 97.8% |
| A21 | 2/LD/T3/3 | 123 | ERS5329264 | 53764366 | 2020-08-24T21:00:00Z | 12 | T3 | 97.5% |
| A22 | 2/LD/T3/4 | 124 | ERS5329265 | 46872910 | 2020-08-24T21:00:00Z | 12 | T3 | 97.8% |
| A23 | 2/LD/T3/5 | 125 | ERS5329266 | 41175448 | 2020-08-24T21:00:00Z | 12 | T3 | 97.8% |
| A24 | 2/LD/T3/6 | 126 | ERS5329267 | 39772946 | 2020-08-24T21:00:00Z | 12 | T3 | 97.6% |
| A25 | 2/LD/T4/1 | 133 | ERS5329268 | 56203190 | 2020-08-25T01:00:00Z | 16 | T4 | 97.6% |
| A26 | 2/LD/T4/2 | 134 | ERS5329269 | 46268272 | 2020-08-25T01:00:00Z | 16 | T4 | 97.4% |
| A27 | 2/LD/T4/3 | 135 | ERS5329270 | 68576730 | 2020-08-25T01:00:00Z | 16 | T4 | 97.0% |
| A28 | 2/LD/T4/4 | 136 | ERS5329271 | 48524028 | 2020-08-25T01:00:00Z | 16 | T4 | 97.3% |
| A29 | 2/LD/T4/5 | 137 | ERS5329272 | 45795270 | 2020-08-25T01:00:00Z | 16 | T4 | 97.4% |
| A30 | 2/LD/T4/6 | 138 | ERS5329273 | 49615352 | 2020-08-25T01:00:00Z | 16 | T4 | 97.7% |
| A31 | 2/LD/T5/1 | 145 | ERS5329274 | 49813802 | 2020-08-25T05:00:00Z | 20 | T5 | 97.9% |
| A32 | 2/LD/T5/2 | 146 | ERS5329275 | 50785920 | 2020-08-25T05:00:00Z | 20 | T5 | 97.7% |
| A33 | 2/LD/T5/3 | 147 | ERS5329276 | 40505336 | 2020-08-25T05:00:00Z | 20 | T5 | 97.6% |
| A34 | 2/LD/T5/4 | 148 | ERS5329277 | 42291530 | 2020-08-25T05:00:00Z | 20 | T5 | 97.9% |
| A35 | 2/LD/T5/5 | 149 | ERS5329278 | 49107588 | 2020-08-25T05:00:00Z | 20 | T5 | 97.7% |
| A36 | 2/LD/T5/6 | 150 | ERS5329279 | 43759616 | 2020-08-25T05:00:00Z | 20 | T5 | 97.6% |
| A37 | 2/LD/T6/1 | 157 | ERS5329280 | 40172870 | 2020-08-25T09:00:00Z | 24 | T6 | 98.0% |
| A38 | 2/LD/T6/2 | 158 | ERS5329281 | 43202190 | 2020-08-25T09:00:00Z | 24 | T6 | 97.6% |
| A39 | 2/LD/T6/3 | 159 | ERS5329282 | 45730066 | 2020-08-25T09:00:00Z | 24 | T6 | 97.8% |
| A40 | 2/LD/T6/4 | 160 | ERS5329283 | 46641334 | 2020-08-25T09:00:00Z | 24 | T6 | 97.7% |
| A41 | 2/LD/T6/5 | 161 | ERS5329284 | 49706180 | 2020-08-25T09:00:00Z | 24 | T6 | 97.4% |
| A42 | 2/LD/T6/6 | 162 | ERS5329285 | 44532910 | 2020-08-25T09:00:00Z | 24 | T6 | 97.7% |

**Table S2.** Clock gene accessions and locations.

| Clock gene | Ensembl Gene_ID | NCBI Accession | NCBI Gene_ID | Chromosome location |
|---|---|---|---|---|
| *clock1a.1* | ENSSSAG00000056165 | XM_014169004.1 | LOC106584123 | ssa23:19987839-20018200 |
| *clock1a.2* | ENSSSAG00000003979 | XM_014122997.1 | LOC106560272 | ssa10:31345809-31398321 |
| *clock2a* | ENSSSAG00000072526 | XM_014137489.1 | LOC106567770 | ssa13:81971735-82044254 |
| *clock2b* | ENSSSAG00000063571 | XM_014196792.1 | LOC106603298 | ssa04:42639734-42675993 |
| *arntl1a.1* | ENSSSAG00000055510 | XM_014148046.1 | LOC106573213 | ssa16:12086304-12095636 |
| *arntl1a.2* | ENSSSAG00000063495 | XM_014125430.1 | LOC106561455 | ssa10:99088660-99104965 |
| *arntl1b.1* | ENSSSAG00000073101 | XM_014175326.1 | LOC106587199 | ssa26:18743473-18765117 |
| *arntl1b.2* | ENSSSAG00000068292 | XM_014126699.1 | LOC106562106 | ssa11:18402203-18411618 |
| *arntl2a* | ENSSSAG00000059155 | XM_014125375.1 | LOC106561420 | ssa10:94715917-94748435 |
| *arntl2b* | ENSSSAG00000008440 | XM_014153577.1 | LOC106576418 | ssa17:47744203-47759123 |
| *arntl2c* | ENSSSAG00000009920 | XM_014208404.1 | LOC106609509 | ssa07:45552387-45568717 |
| *per1a.1* | ENSSSAG00000077896 | XM_014196530.1 | LOC106603198 | ssa04:38522751-38532432 |
| *per1a.2* | ENSSSAG00000063164 | XM_014128896.1 | LOC106563389 | ssa11:73693186-73705207 |
| *per1b* | ENSSSAG00000002804 | XM_014206471.1 | LOC106608520 | ssa07:4731276-4739498 |
| *per2a* | ENSSSAG00000076981 | XM_014139595.1 | LOC100301980 | ssa14:16742620-16769326 |
| *per2b* | ENSSSAG00000001704 | XM_014190838.1 | LOC106599551 | ssa03:17486279-17517618 |
| *per3* | ENSSSAG00000077674 | XM_014166967.1 | LOC106583133 | ssa22:25358497-25382546 |
| *cry1a.1* | ENSSSAG00000077590 | XM_014148221.1 | LOC106573292 | ssa16:15906400-15976136 |
| *cry1a.2* | ENSSSAG00000071693 | XM_014125353.1 | LOC106561407 | ssa10:95086446-95106453 |
| *cry2* | ENSSSAG00000073888 | XM_014125728.1 | LOC106561609 | ssa10:105488616-105553942 |
| *cry3b.1* | ENSSSAG00000081201 | XM_014131850.1 | LOC106565112 | ssa12:46213293-46226192 |
| *cry3b.2* | ENSSSAG00000081121 | XM_014167724.1 | LOC106583488 | ssa22:40956787-40966869 |
| *cry4* | ENSSSAG00000069768 | XM_014176427.1 | LOC106587781 | ssa26:36623393-36647646 |
| *cry5* | ENSSSAG00000079794 | XM_014138346.1 | LOC106568206 | ssa13:86762607-86774549 |
| *nr1d1a* | ENSSSAG00000003153 | XM_014204497.1 | LOC106607496 | ssa06:40433770-40444554 |
| *nr1d1b* | ENSSSAG00000007825 | XM_014192527.1 | LOC106600825 | ssa03:57681841-57689097 |
| *nr1d2a.1* | ENSSSAG00000037545 | XM_014164862.1 | LOC106582110 | ssa02:25864163-25873212 |
| *nr1d2a.2* | ENSSSAG00000005012 | XM_014200712.1 | LOC100136378 | ssa05:54986045-54995902 |
| *nr1d2b.1* | ENSSSAG00000056065 | XM_014142438.1 | LOC106570286 | ssa14:68249965-68261020 |
| *nr1d2b.2* | ENSSSAG00000054663 | XM_014177878.1 | LOC106588668 | ssa27:19843632-19851666 |
| *nr1d4a.1* | ENSSSAG00000065806 | XM_014147027.1 | LOC106572653 | ssa15:88787769-88813331 |
| *nr1d4a.2* | ENSSSAG00000077410 | XM_014135912.1 | LOC106567064 | ssa13:35389895-35404997 |
| *nr1d4b.1* | ENSSSAG00000081117 | XM_014167340.1 | LOC106583303 | ssa22:32346247-32358789 |
| *nr1d4b.2* | ENSSSAG00000067342 | XM_014132672.1 | LOC106565463 | ssa12:54465885-54474575 |
| *roraa.1* | ENSSSAG00000066827 | XM_014148628.1 | LOC106573506 | ssa16:27588967-27706002 |
| *roraa.2* | ENSSSAG00000074849 | XM_014124804.1 | LOC106561150 | ssa10:82919138-82990987 |
| *rorb.1* | ENSSSAG00000069458 | XM_014160683.1 | LOC106580068 | ssa20:14747696-14776762 |
| *rorb.2* | ENSSSAG00000065430 | XM_014172220.1 | LOC106585703 | ssa24:30374236-30414448 |
| *rorca.1* | ENSSSAG00000073620 | XM_014169656.1 | LOC106584392 | ssa23:30486289-30543797 |
| *rorca.2* | ENSSSAG00000068177 | XM_014216453.1 | LOC106613816 | ssa10:21617008-21834653 |
| *rorcb.1* | ENSSSAG00000079594 | XM_014140953.1 | LOC106569518 | ssa14:38121993-38144650 |
| *rorcb.2* | ENSSSAG00000080721 | XM_014191359.1 | LOC106600026 | ssa03:37981172-38001032 |
| *csnk1db.1* | ENSSSAG00000068205 | XM_014179815.1 | LOC106589616 | ssa28:16447101-16461477 |
| *csnk1db.2* | ENSSSAG00000003495 | XM_014209485.1 | LOC106610228 | ssa01:77662396-77680525 |
| *csnk1ea* | ENSSSAG00000001530 | XM_014193266.1 | LOC106601228 | ssa03:65302128-65311565 |
| *csnk1eb* | ENSSSAG00000068400 | NM_001140384.1 | LOC100195355 | ssa06:33906202-33916152 |
| *csnk1ec* | ENSSSAG00000015336 | XM_014179651.1 | LOC106589542 | ssa28:14144088-14151122 |
| *csnk1ed* | ENSSSAG00000003438 | XM_014208340.1 | LOC106609470 | ssa01:74533732-74548069 |

**Table S3.** Number of significant genes: Rhythmic (RAIN) or circadian (JTK) depending on the P-value (FDR adjusted) or relative Amplitude thresholds used. All dataset and clock genes only (in bracket).

| | **RAIN** | **JTK** | **overlap** |
|---|---|---|---|
| adjP-value $< 0.05$ | | | |
| rAmp $\geq 0$ | 12,322 (32) | 6,485 (22) | 6,427 (22) |
| rAmp $\geq 5\%$ | 9,818 (25) | 5,954 (21) | 5,896 (21) |
| rAmp $\geq 10\%$ | 5,529 (20) | 3,678 (18) | 3,621 (18) |
| rAmp $\geq 15\%$ | 2,906 (8) | 2,007 (8) | 1,959 (8) |
| adjP-value $< 0.01$ | | | |
| rAmp $\geq 0$ | 9,064 (27) | 3,786 (15) | 3,760 (15) |
| rAmp $\geq 5\%$ | 7,413 (24) | 3,627 (15) | 3,602 (15) |
| rAmp $\geq 10\%$ | 4,234 (19) | 2,374 (14) | 2,349 (14) |
| rAmp $\geq 15\%$ | 2,206 (7) | 1,264 (6) | 1,241 (6) |
| adjP-value $< 0.001$ | | | |
| rAmp $\geq 0$ | 5,815 (22) | 1,721 (11) | 1,717 (11) |
| rAmp $\geq 5\%$ | 4,911 (21) | 1,702 (11) | 1,698 (11) |
| rAmp $\geq 10\%$ | **2,864 (16)** | **1,215 (11)** | **1,211 (11)** |
| rAmp $\geq 15\%$ | 1,470 (6) | 648 (5) | 644 (5) |
